# Local reflects global: Life-stage dependent changes in the phenology of coastal habitat use by North Sea herring

**DOI:** 10.1101/2023.11.23.568404

**Authors:** Mark Rademaker, Myron A. Peck, Anieke van Leeuwen

## Abstract

Climate warming is affecting the suitability and utilisation of coastal habitats by marine fishes around the world. Phenological changes are an important indicator of population responses to climate-induced changes but remain difficult to detect in marine fish populations. The design of large-scale monitoring surveys does not allow fine-grained temporal inference of population responses, while the responses of ecologically and economically important species groups such as small pelagic fish are particularly sensitive to temporal resolution. Here, we use the longest, highest-resolution time series of species composition and abundance of marine fishes in northern Europe to detect possible phenological shifts in the small pelagic North Sea herring. We detect a clear forward temporal shift in the phenology of nearshore habitat use by small juvenile North Sea herring. This forward shift can best be explained by changes in water temperatures in the North Sea. We find that reducing the temporal resolution of our data to reflect the resolution typical of larger surveys makes it difficult to detect phenological shifts and drastically reduces the effect sizes of environmental covariates such as seawater temperature. Our study therefore shows how local, long-term, high-resolution time series of fish catches are essential to understand the general phenological responses of marine fishes to climate warming and to define ecological indicators of system-level changes.

## Introduction

Climate warming has caused well-documented shifts in the distribution of species in terrestrial and aquatic ecosystems [1] including changes in the dynamics of fish populations across the globe [2, 3, 4]. In several coastal zones, phenological shifts in the occurrence of adult and larval fish have also been observed [5, 6]. In parallel, industrial fishing has altered the abundance, structure, and reproductive characteristics of marine fish populations [7, 8, 9, 10]. As opposed to large-scale effects on populations that may be readily detectable, understanding the role of external environmental processes on local or regional population dynamics is more challenging [11]. Small pelagic fish species pose an especially complex case, as their populations can exhibit both ‘volatile’ short-term dynamics in response to local or regional conditions, and long-term ‘stable’ cyclicity due to larger scale climate events [12, 13]. Furthermore, many species exhibit ontogenetic shifts in habitats, spending part of their life cycle in coastal or estuarine zones, which may constrain the ability of species to cope with warming [14]. The importance of external environmental processes can, therefore, be life stage-specific [15, 16]. The local dynamics of small pelagic fish are an essential component to better understand potential changes in areas historically important to the life cycle dynamics of species, and to examine whether the phenology in the use of important habitats may have shifted over time.

The Atlantic herring (*Clupea harengus*) is a common, ecologically and commercially important small pelagic fish species in areas of the North Atlantic such as the North Sea [17, 18] where the population is composed of separate autumn- and winter-spawning stocks with life stage-specific distributions [19]. In the southern North Sea, Atlantic herring is also commonly observed in the Wadden Sea, which with its shallow coastal waters serves as a spawning, nursery, and feeding ground for many North Sea fish species [20]. However, much is unknown about the habitat utilization of the Wadden Sea by Atlantic herring. This is primarily because herring and other small pelagic fishes cannot be properly sampled in the demersal trawl surveys used to annually monitor fish populations in these near-shore waters [21]. Furthermore, the coarse temporal resolution of these large-scale surveys does not allow inferences on seasonally important factors. Limited temporal resolution also prevents the detection of potential phenological shifts that might have occurred during the past few decades. This limits our understanding of the external processes governing the local dynamics of herring and other fish moving between shallow coastal nursery and feeding grounds of the Wadden Sea and offshore waters of the North Sea. Local survey efforts conducted at high temporal resolution are needed to help fill such important gaps in knowledge on potential climate-driven changes in phenology and habitat utilization.

Survey efforts with sufficiently high temporal resolution to examine climate-driven changes in phenology are rare in the marine environment, particularly those with a historical coverage allowing comparisons over multiple decades. Although long term high temporal resolution time-series exist for lower trophic levels such as plankton [22], they are particularly rare for higher trophic levels such as fish. However, the Royal Netherlands Institute for Sea Research (NIOZ) has consistently used a kom-fyke to collect standardized catch data of marine fishes moving between the North Sea and Western Wadden Sea through the Marsdiep tidal race since the 1960’s [23] (Fig. 1). This survey is unique in northern Europe due to its daily temporal resolution. Previous analyses of these data have identified phenological shifts such as changes in the day of first occurrence, peak occurrence, and last occurrence of members of the Wadden Sea fish community over the past five decades [24]. Given its daily resolution, this time series allows a finer-scale exploration of changes in the phenology of species such as Atlantic herring. From a methodological standpoint, the preselection of specific days (first occurrence, peak occurrence, last occurrence) as a phenological yardstick for peak migration may obscure shifts in species such as herring that, due to their schooling and flexible foraging behaviour, exhibit marked daily variation in habitat use or occupancy. Furthermore, it remains unknown if and how potential shifts in phenology differ among young juvenile and adult herring. More detailed phenological analysis would reveal if local dynamics in coastal habitat use by North Sea herring align with the larger regional and global population responses to climate-driven warming observed in marine fish.

**Figure 1.**
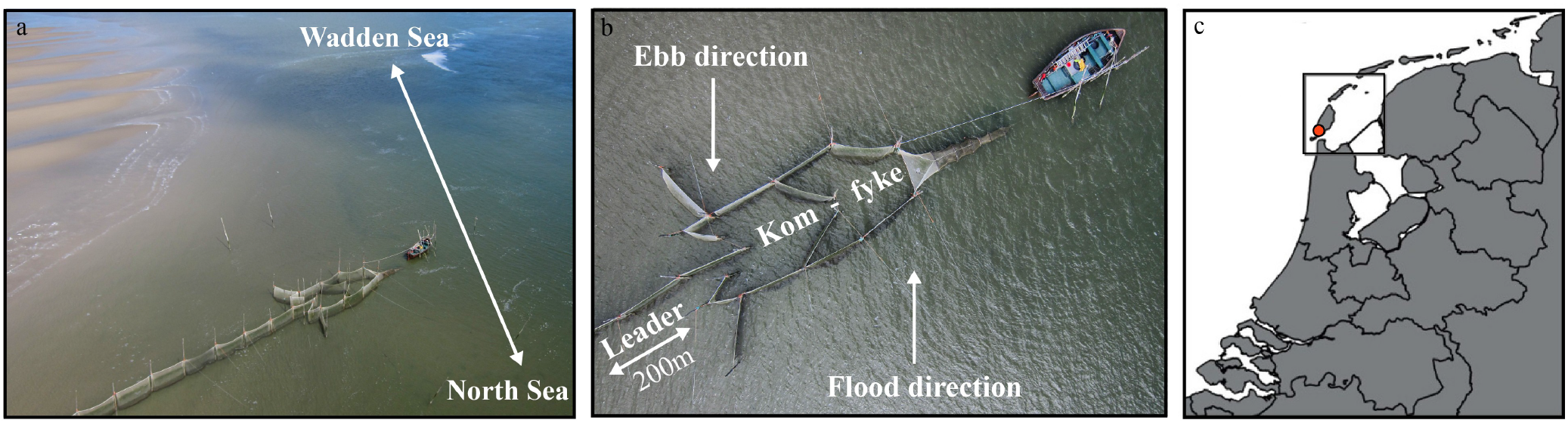
Set-up of the NIOZ kom-fyke (a,b) and its geographic position on the border between the North Sea and Wadden Sea (c). Images a and b used with permission from van Walraven (personal communication).

Here, we examined potential life-stage specific changes in the population dynamics of Atlantic herring moving between the North Sea and Wadden Sea using long-term time series of weekly standardized herring catches (1981-2021) collected by the NIOZ kom-fyke. We used a generalized additive modelling (GAM) approach to decompose the overall trend in weekly standardized herring catches into distinct annual, seasonal, and environmental signals. These results allowed us to identify temporal trends in herring abundance, body size, and reproductive status in the Western Dutch Wadden Sea. We also examined potential changes in the seasonal trends in herring abundance over the past four decades, i.e. phenological shifts. Finally, we checked the sensitivity of our outcomes with respect to changes in the sampling design of the Kom-fyke by re-running our analysis with different sampling frequencies. Our analyses on the longest, most high-resolution time series for marine fish in Northern Europe will advance understanding of the factors determining the movement and phenological shifts of herring and potentially other temperate small pelagic fishes from offshore to nearshore waters. Our work underscores the importance of maintaining highly temporally resolved, long-term ecological time series for examining the dynamics of climate-driven phenological shifts.

## Materials and Methods

To examine phenological changes in North Sea herring and the factors contributing to potential changes in these dynamics, we analysed the number of herring in daily spring and autumn catches from 1982 - 2021. We also included information on fish life-stage, based on dissections of herring captured in the same program. Below we describe the source of the data and steps in data preparation and analysis in detail.

### Data sampling

#### Fish catches

We used catches in the long-term time-series data from the NIOZ kom-fyke program. The leader net of the kom-fyke extends 200 meters from the shoreline into the subtidal zone where two chambers with a mesh-size of 10 x10 mm collect fishes [23]. Additional details on the kom-fyke netting and other gear and program specifications are provided by van der Veer et al. [25] . The kom-fyke is emptied daily during two annual sampling periods that run approximately from the end of March through the end of June, and from the start of September through the beginning of November. The timing of this monitoring coincides with the well-known seasonal ingress of fish from the North Sea into the Wadden Sea in spring, and the emigration of young-of-the-year fish from the Wadden Sea to the North Sea in autumn. Kom-fyke sampling is discontinued in high summer due to net fouling by macro-algae blooms. Sampling does not occur in winter due to increased risks to equipment and personnel posed by storms and ice floes. Catches were sorted according to species and the total length (TL, ± 0.5 cm) of each individual was measured. If herring catches were too large to count all specimens individually, a well-mixed, volume-based subsample was taken, to ensure all different size classes remained proportionally represented. For the historical developments of the fyke program and sample counting see van der Veer [23].

#### Abiotic factors

We collected data on local Wadden Sea surface water temperature, regional North Sea surface water temperatures, lunar illumination, and local tidal range as abiotic explanatory variables. Local Wadden Sea surface water temperatures have been measured continuously by the NIOZ Jetty monitoring setup [26], located approximately 500 meters from the fyke (Fig. 1). Regional North Sea surface water temperatures and tidal ranges near the fyke (Den Helder) were collected from publicly accessible weather buoy and acoustic sensor data [27]. We calculated the maximum possible lunar illumination received for each sampling date using the Lunar package in R [28].

### Data preparation

#### Phenology data

We defined a time window when daily sampling data were available from both the fyke sampling and abiotic factors for a 39-year period (1982-2021). We performed additional filter operations on the dataset to minimize any bias due to data measurement or entry errors, as well as potential overestimation of catches due to weather events preventing the daily emptying of the fyke. We only included samples with >12 and <48 hours of fishing effort, and samples with reported body sizes falling within the species limits known from literature. The catch data were standardized by taking the weekly summed catch per unit effort (CPUE), and the weekly mean of the abiotic factors. We chose weekly over daily CPUE to correct for a lack of fishing on some days. The final time-series consisted of 934 sampled weeks of log-transformed herring catches and abiotic factors spread over 39 years. Resolution in the time series varied slightly during two periods. First, there were no catch data from early 2020 when the covid pandemic prohibited fieldwork. Second, the temporal resolution in regional North Sea water temperatures was lower for certain weeks in the 1980’s compared to 1990-2021. Therefore, we ran our model multiple times, excluding and including these parts of the time series, to examine qualitative, and quantitative changes in model outcomes. We found that only the effect of tidal range changed depending on the inclusion of these time periods, but the effect of all other covariates remained the same. We therefore chose to include these periods in the time-series and excluded the tidal range effect from detailed interpretation.

#### Dissection data

We used dissection data performed on a subset of weekly catches from 2005-2021 to assess trends in the reproductive status of herring in the kom-fyke (N=480). The reproductive status of individuals was determined based on a six-point scale of gonadal ripening (Supplementary Table 1). We used this information to determine the length at which 50% of individuals had reached maturity using logistic regression (L_m_50) (Supplementary Figure 1). The abundance data were then divided into juvenile (TL < cm) and adult (TL > cm) categories to examine how the relative contribution of these two life stages to catches might have shifted over time.

### Data analysis

We used generalized additive models (GAMs, [29]) to examine temporal changes in the local population dynamics of herring moving between the Wadden Sea and North Sea in relation to abiotic factors. We fitted our additive models using the function gam from the package mgcv [30]. Our base-model included North Sea and Wadden Sea water temperatures, lunar illumination, tidal range, week, year, and the interaction between week and year as covariates, next to a temporal autocorrelation component *ρ* (eqn. 1).

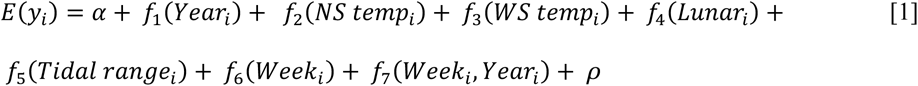

Where smooths *f*_1_ – *f*_6_ represent the main effects of year, North Sea and Wadden Sea water temperatures, lunar illumination, tidal ranges, and week respectively. The smooth *f*_7_ is a tensor interaction product that models how the seasonal effect of week on herring catches varies over the years. A tensor interaction allows assessment of the separate singular effects of two variables versus their interaction effect, which is why we chose this type of interaction term. We used the automated variable selection procedure developed by Marra and Wood to identify significant model terms [31]. Next to this, we checked for model fit by examining model convergence and residual diagnostic tables and plots (Quantile-Quantile, density distribution, response vs. fitted values, and autocorrelation) using the *gam.check()* and *resid.check()* functionalities in the R package mgcv and itsadug respectively [30, 32]. The base model converged, explained 43.5% of variation, 45% of null deviance, and had good residual fit diagnostics (Table 1, Supplementary Figure 2,3). We checked for potential extrapolation artifacts in seasonal trends due to the temporal gap between fishing seasons within a year by running the same model on subsets of spring data only (R^2^ = 29.2%; Deviance = 31.2%) and fall data only (R^2^ = 33.1%; Deviance = 34.8%). The predicted week trend and the interaction between week and year remained qualitatively similar between these models and the full-data model (Supplementary Figure 4, 5).

**Table 1.** Smoother fit diagnostics. Low p-values (k-index <1) can indicate the number of knots used in constructing the smoother is too low. Except for S(Year), all smoothing terms in the model had good fit. We varied the number of knots in S(Year), but found no difference in smoother shape or fit, and therefore kept the automatically selected smooth with 9 knots in the model.

| Smooth | Spline | Knots | Edf. | K-index | p-value |
| --- | --- | --- | --- | --- | --- |
| S(Year) | Thin plate | 9 | 2.326 | 0.60 | < 0.001 |
| S(NS temp) | Thin plate | 9 | 0.747 | 0.96 | 0.13 |
| S(WS temp) | Thin plate | 9 | 1.286 | 1.01 | 0.56 |
| S(Tidal range) | Thin plate | 9 | 0.403 | 1.05 | 0.89 |
| S(Lunar) | Thin plate | 9 | 2.038 | 1.02 | 0.69 |
| S(Week) | Cyclic cubic | 8 | 6.102 | 1.06 | 0.93 |
| S(Week, Year) | Cyclic cubic, Thin plate | 12 | 2.12 | 1.04 | < 0.001 |

Next, we attempted to expand our base model to examine the role of body size in driving the phenological trends by including herring size-class in cm’s as an additional covariate. The expanded model with continuous body size converged, and explained 47% of variation, implying that body sizes carry additional relevant information. However, the model also suffered from considerably worse fit, especially heteroskedasticity. Residual plots indicated that the model could not adequately predict small catch numbers close to zero, systematically overestimating them. This is due to zero-inflation when accounting for continuous size classes, i.e. for most size classes catches are zero. To address this issue, we first tried fitting a zero-inflated negative binomial, and a zero-inflated poisson distribution to untransformed catch data. However, due to the large range and variation in catches (0-5000), choosing a zero-inflated poisson or negative binomial distributions on untransformed catches did not improve fit. Fit could be improved when only ‘average’ catches were included. However, the high ‘outlier’ catches are of particular interest in this study as they represent peak migration points. We therefore chose to examine how the role of body size and life stage have shifted over time by visually inspecting ridgeline plots, rather than including size class as a covariate in the model.

Finally, we examined how the modelled effects change with respect to monitoring design by rerunning the model with lower sampling frequencies (single fishing day per week, single fishing day per two weeks, single fishing day per month). We used these frequencies to have an approximate qualitative comparison to the outcomes expected in large fisheries surveys with limited temporal resolution. For the single sample per week case, we used the first fishing day of every week, for the biweekly case we used the first fishing day of every odd week, and for the monthly sample we used the first fishing day of the month. We examined model fit and summary statistics and extracted the predicted partial effects for each covariate to examine how both the significance and the effect size changed under each sampling regime.

## Results

The abundance of herring fluctuated strongly during the 1982 -2021 period (Fig. 2a; Mean weekly standardized catch: 3 198 ± 10 501 individuals, Median weekly standardized catch: 356 individuals). Our additive model reflected these fluctuations, predicting a large seasonal component that matched the peak-to-peak periodicity, but not the amplitude, of observed herring abundance. The model, therefore, underestimated the absolute values of the minima and maxima in herring catches. The fitted trend could be decomposed into significant partial smoother effects of year, week, lunar illumination, tidal range, and regional North Sea water temperatures (Table 2).

**Figure 2.**
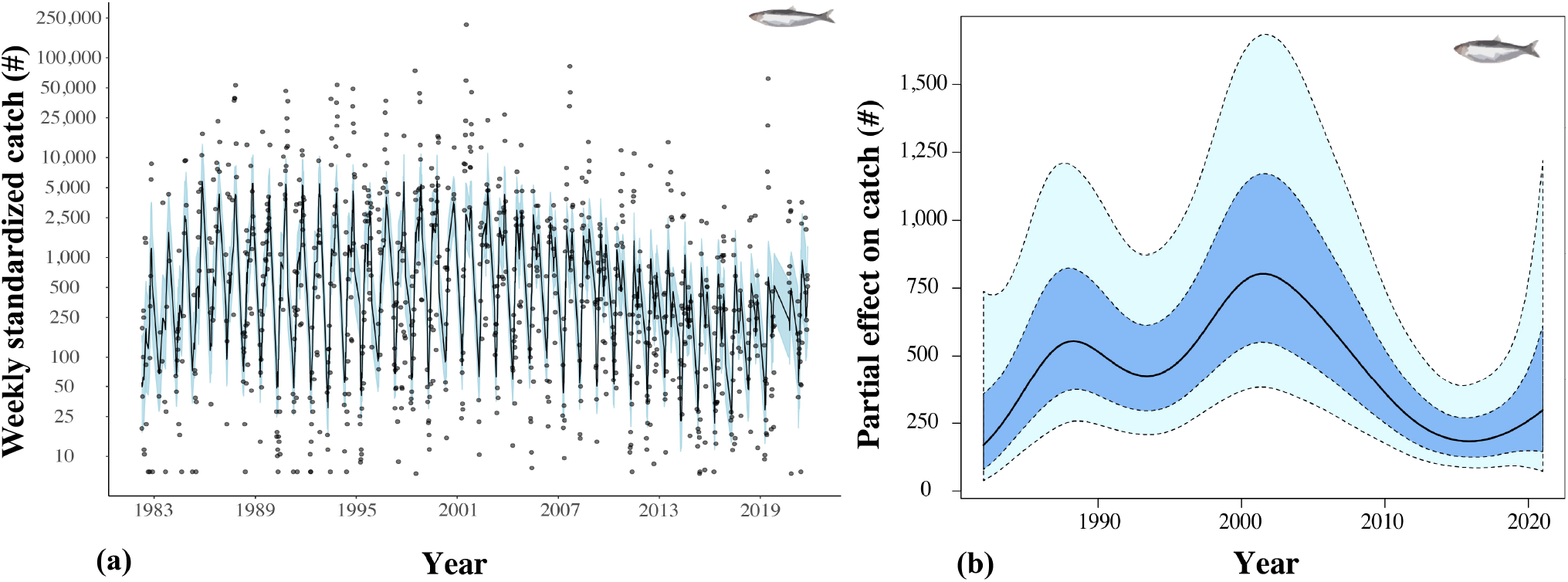
**(a)** Fitted additive model predictions of weekly standardized herring catches ±95% CI in the NIOZ kom-fyke from 1982 - 2021. Grey dots represent the observed values. The gap in early 2020 is due to covid restrictions prohibiting fieldwork. Rerunning the model excluding the gap did not alter the shape or significance of individual smooth terms in the generalized additive model **(b)** Partial effect of year on weekly standardized herring catches. Shaded dark blue area represents the point-wise standard errors, and the shaded light blue area the point-wise 95% CI.

**Table 2.**
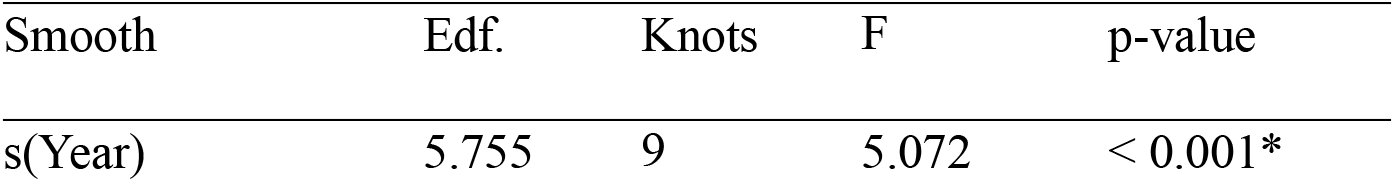
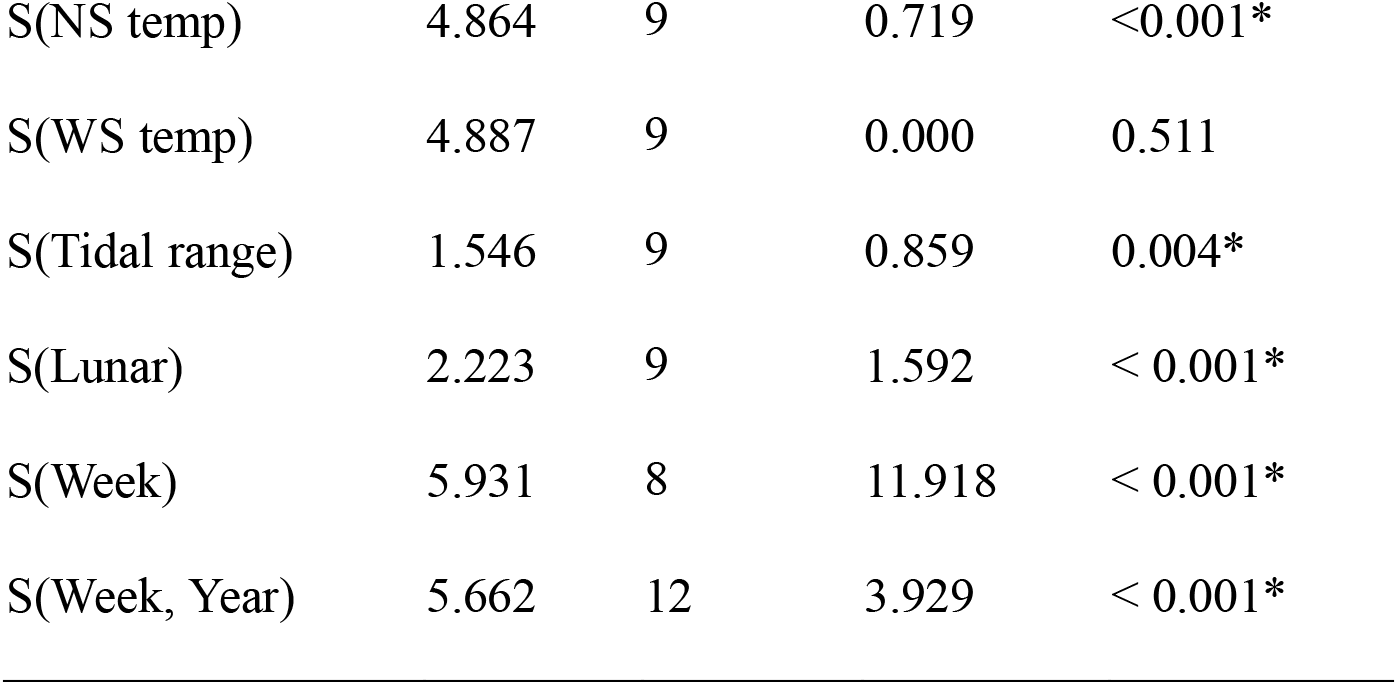
Approximate significance of covariate smoother terms in the additive model (R2 = 43.5%, Deviance = 45%). Significance of a covariate smoother indicates that the null-hypothesis is rejected, which means that the partial additive effect of a covariate upon weekly standardized catches cannot be modelled using a flat line. The ecological significance of the effect must be interpreted based on the effect size.

### Partial effects

The partial effect of year identified strong periods of increase and decrease in predicted herring catches during the past four decades (Fig. 2b). Weekly standardized catches were predicted to be lowest at the start of the time series in 1982 with 169 individuals (95% CI (38, 738). Subsequently, there was a period of gradual increase in catches, up to 803 individuals, 95% CI (383, 1 684) in 2001. However, this period was followed by a strong decrease back to low levels of 183 individuals, 95% CI (86, 391) in 2015, and 298 individuals, 95% CI (73, 1 219) in 2021. The narrowing and decrease in upper and lower confidence bounds during the period of decline in the last two decades indicate a general decrease in peak weekly standardized catches in individual years during this time period.

We find a strong seasonal signal in herring catches depending on the week sampled within the year (Fig. 3a). Predicted catches differed by up to two orders of magnitude between weeks 10-20 (∼ 25 individuals, 95% CI (6, 110)) and weeks 30-40 (∼ 1 186 individuals, 95% CI (261, 5 379)). We also observed a forward temporal shift in the seasonal trend over time (Fig. 3b), with a relative increase in predicted catches in spring and summer (∼ weeks 16-30), and a relative decrease in predicted catches in late fall (∼ weeks 40-50).

**Figure 3.**
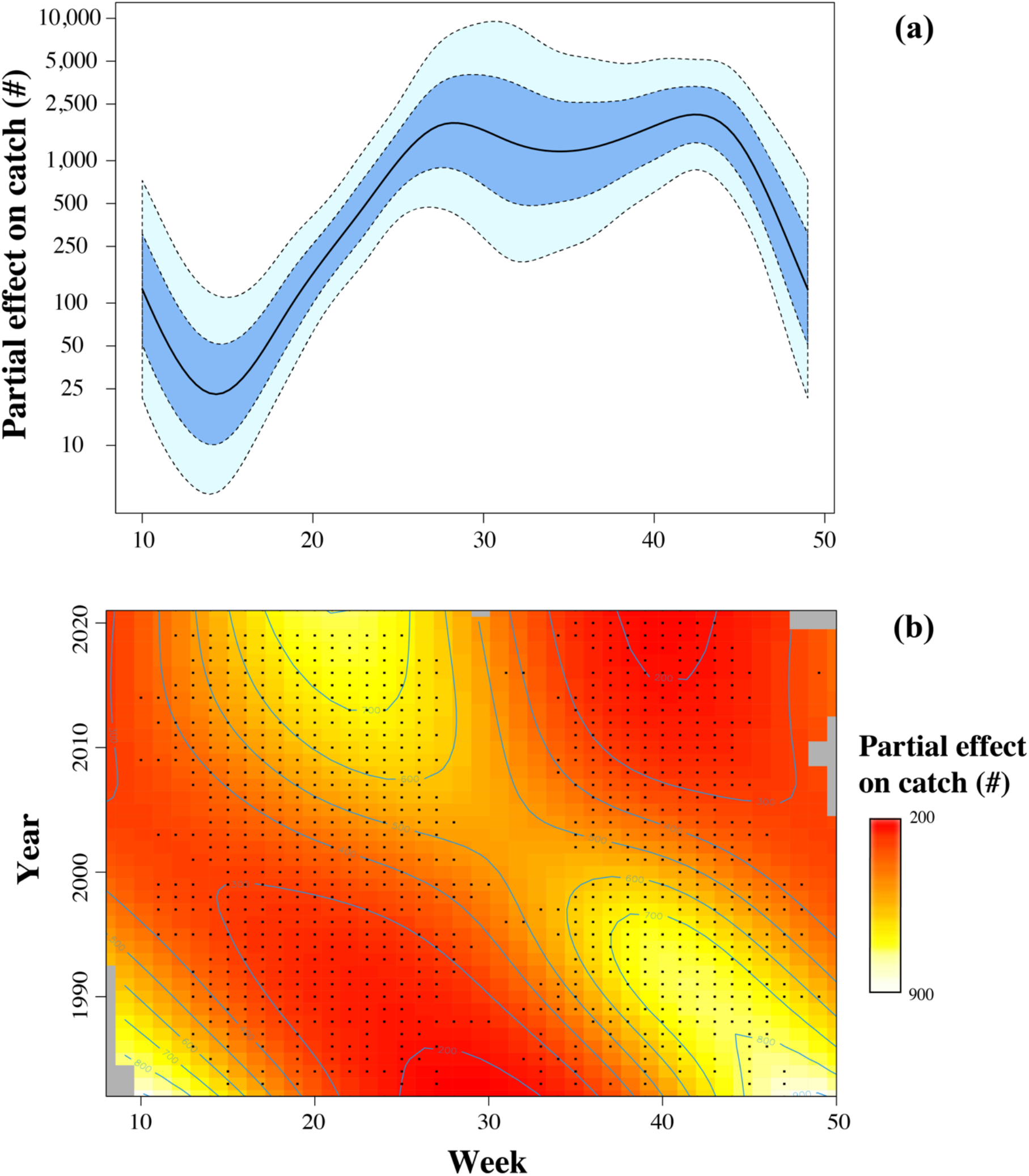
**(a)** Partial effect of week number on weekly standardized herring catches. Shaded dark blue area represents the point-wise standard errors, and the shaded light blue area the pointwise 95% CI. **(b)** Partial interaction effect of week and year on standardized herring catches. Over time there has been a relative increase in catches early in the year (weeks 15 – 22) and a relative decrease in catches late in the year (weeks 35 – 45). The color bar shows the magnitude of the effect. The sampling effort is divided into two catch seasons per year (spring and fall) represented with black dots. Consistency of the effect with respect to sampling season is shown in Supplementary Figure 4 & 5.

Regional water temperatures in the North Sea were found to be more important for changes in the abundance of herring than local water temperatures in the Wadden Sea. Relatively cold North Sea water temperature (5-10*^◦^*C) lead to approximately five-fold increases in predicted catches (∼ 1 554 individuals, 95% CI (267, 9 023)) compared to those expected under the long-term mean annual water temperature of 12.8 *^◦^*C (348, 95% CI (171, 709)) (Supplementary Figure 6). Lunar illumination had a much smaller effect on changes in herring abundance. Periods of full moon and the change to new moon increased weekly standardized catches by ∼ 200 individuals compared to catches outside these periods (∼ 517 vs. 329 individuals; Supplementary Figure 7). The effect of tidal range varied between periods of high and low tidal ranges, but we found this effect to be inconsistent with respect to time-series resolution, whereas all other partial effects were consistent. We, therefore, exclude tidal range from detailed interpretation.

### Sampling frequency

We find that moving from weekly standardized catches based on daily sampling to standardized catches based on a single fishing day every week, biweekly, or per month, significantly changed effect sizes and caused poor model fits due to heteroskedasticity (Table 3; Supplemental Figures 8-10). The effect of North Sea water temperature was no longer found when sampling frequency was decreased. The lunar effect disappears when samples were taken biweekly or monthly. In contrast, the significant effect of year, season, and changes in the seasonal effect over years remained present for all sampling frequencies. However, the effect sizes were reduced by almost a hundred-fold with decreased sampling frequency. Visual inspection indicated that the resulting smoother plots were qualitatively similar in the year and seasonal trend between sampling frequency designs (Supplementary Figure 11, 12), but there were large differences in the plots of changing seasonal trends over time, complicating the interpretation of phenological shifts at reduced sampling frequencies (Supplementary Figure 13). Only in the monthly sampling case is a phenological shift visible but the effect size of the change in standardized catches over time was negligible.

**Table 3.**
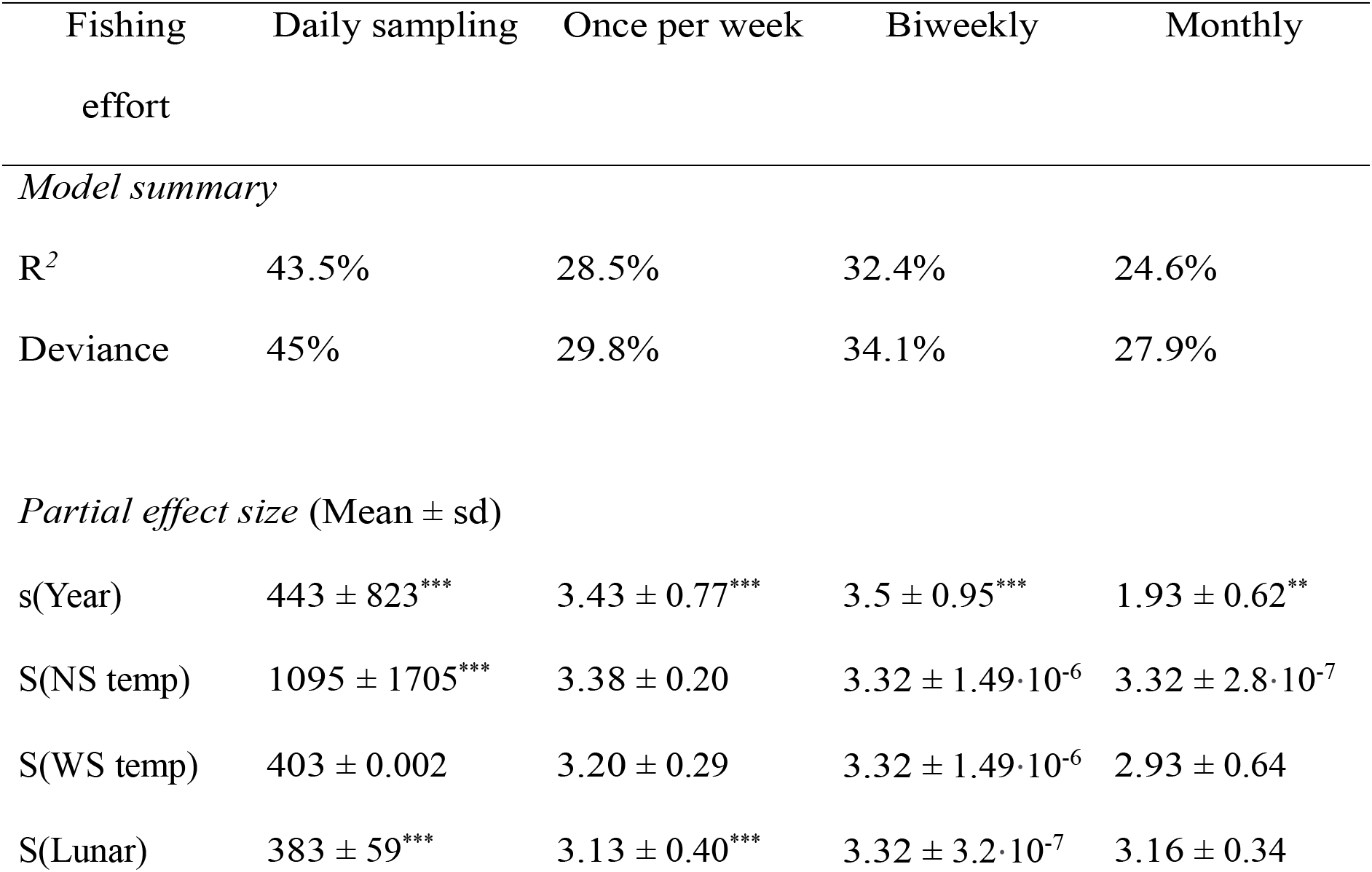
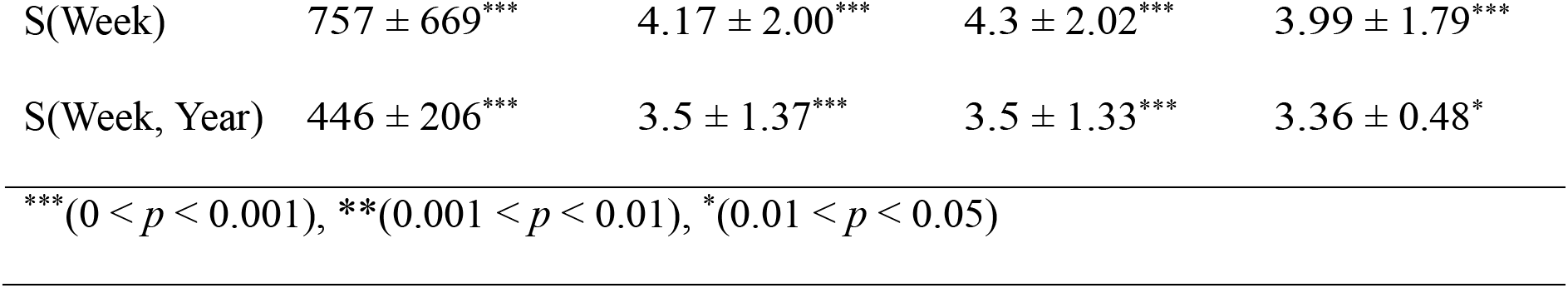
Comparison of model performance and approximate covariate effect sizes under different sampling frequencies. Approximate effect sizes were calculated by extracting the predicted partial effect sizes for each individual covariate from the model and taking the mean and the standard deviation. A mean with a high standard deviation indicates a non-constant and likely significant effect size of the covariate, while a mean with a low standard deviation indicates the fitted effect can be represented by a flat line and the covariate is therefore likely unsignificant.

| Fishing effort | Daily sampling | Once per week | Biweekly | Monthly |
| --- | --- | --- | --- | --- |
| *Model summary* |  |  |  |  |
| R<sup>2</sup> | 43.5% | 28.5% | 32.4% | 24.6% |
| Deviance | 45% | 29.8% | 34.1% | 27.9% |
| *Partial effect size (Mean ± sd)* |  |  |  |  |
| s(Year) | 443 ± 823*** | 3.43 ± 0.77*** | 3.5 ± 0.95*** | 1.93 ± 0.62** |
| S(NS temp) | 1095 ± 1705*** | 3.38 ± 0.20 | 3.32 ± 1.49·10<sup>-6</sup> | 3.32 ± 2.8·10<sup>-7</sup> |
| S(WS temp) | 403 ± 0.002 | 3.20 ± 0.29 | 3.32 ± 1.49·10<sup>-6</sup> | 2.93 ± 0.64 |
| S(Lunar) | 383 ± 59*** | 3.13 ± 0.40*** | 3.32 ± 3.2·10<sup>-7</sup> | 3.16 ± 0.34 |
| S(Week) | 757 ± 669<sup>***</sup> | 4.17 ± 2.00<sup>***</sup> | 4.3 ± 2.02<sup>***</sup> | 3.99 ± 1.79<sup>***</sup> |
| S(Week, Year) | 446 ± 206<sup>***</sup> | 3.5 ± 1.37<sup>***</sup> | 3.5 ± 1.33<sup>***</sup> | 3.36 ± 0.48<sup>*</sup> |

<sup>\*\*\*</sup>(0 < *p* < 0.001), <sup>\*\*</sup>(0.001 < *p* < 0.01), <sup>\*</sup>(0.01 < *p* < 0.05)

### Changes in body size

The relative contribution of size classes remained similar over time but differed markedly between seasons (Fig. 4). In both the spring and autumn seasons, catches were dominated by juveniles between 5 - 15 cm in TL. Although larger adult herring (20 - 30 cm TL) were also abundant in samples collected in spring and summer, they were very rare in autumn and winter samples. Furthermore, compared to the 1980s, smaller fish have become more abundant with smaller minimum size classes observed in the most recent decade. The size class distributions provided evidence that small juvenile herring drive the observed changes in the kom-fyke time series as opposed to changes in catches of adult individuals (Fig. 4).

**Figure 4.**
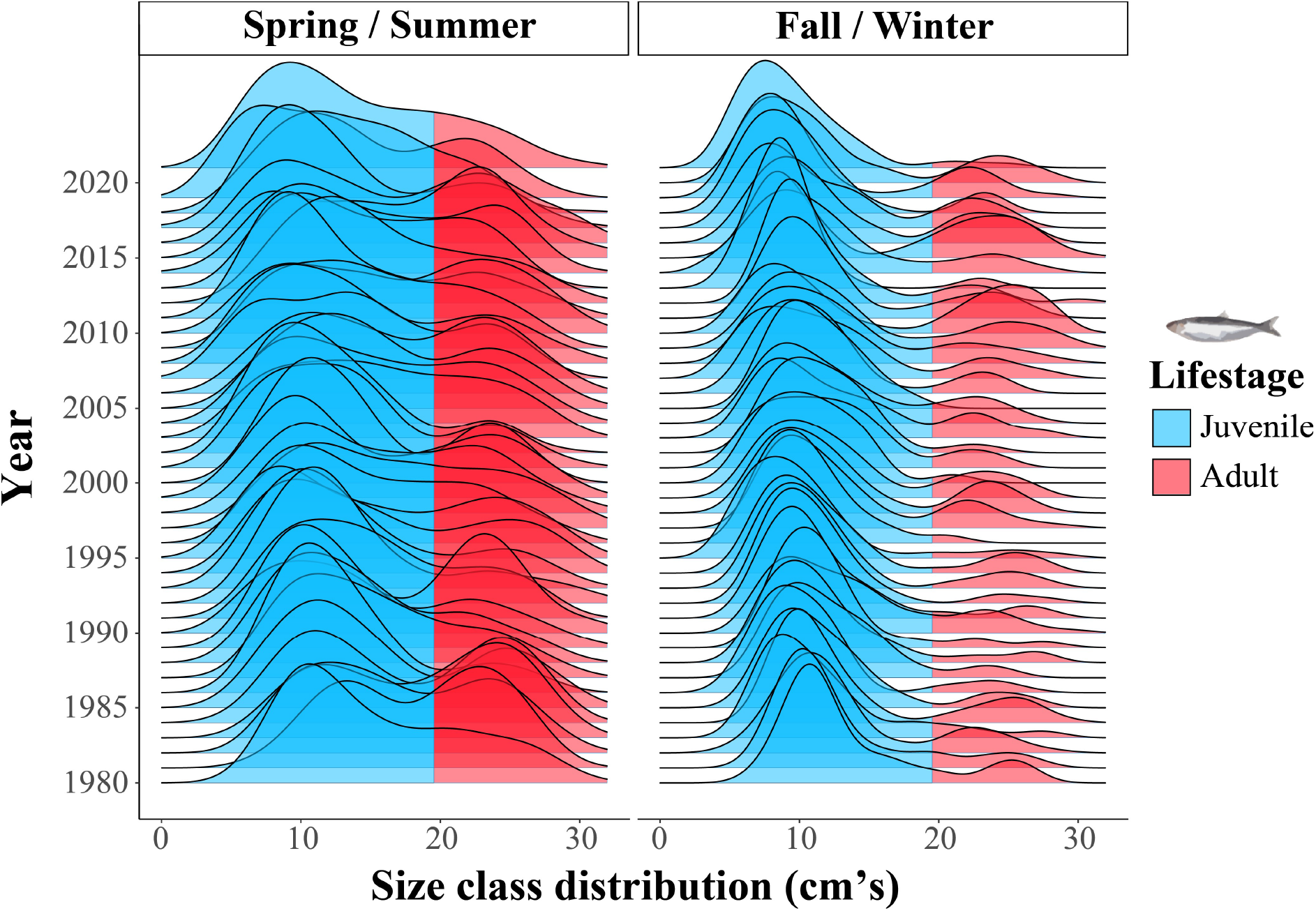
Annual and seasonal distributions of size classes in standardized herring catches from the kom-fyke. Colors are illustrative for the L_m_50 maturation size of dissected individuals (19.5 cm, Supplementary Figure 1), although there will be small year-to-year differences in maturation size.

## Discussion

Our analyses of the longest, highest-resolution time-series of marine fish species composition and abundance in Northern Europe detects phenological shifts in North Sea herring. Phenological shifts are challenging to detect in ecosystems but can indicate important climate-driven changes in the suitability and use of different habitats [33, 34]. Changes in the spatial coverage and timing of survey efforts between years and geographic heterogeneity between survey locations can all confound the detection and interpretation of phenological changes [35, 36, 37]. In marine fish populations, the detection of phenological shifts is further complicated by tradeoffs in the spatial versus temporal sampling resolution of survey designs and the specificity of sampling gear [35]. The longest and spatially largest marine fish surveys, such as bottom trawl surveys in the northeast and northwest that target benthic species, are performed either annually or quarterly and follow a random stratified sampling design [38]. These surveys, designed to monitor the abundance and biomass of commercially important marine fish, have proven to be powerful in detecting broad horizontal (longitudinal or latitudinal) or depth shifts of fish in response to climate-driven warming [39, 40]. However, these survey designs possess several factors that confound phenological analyses, namely their low temporal resolution, interannual variation in sampling locations, changing spatial coverage, and bias in species catchability.

As opposed to long-term quarterly or annual broad-scale surveys, long-term ‘local’ monitoring surveys with high temporal resolution have been very important in detecting climate-driven changes in the phenology and coastal habitat use of fish communities, e.g. in Narragansett Bay, RI (USA) [5]. It remains challenging, however, to document phenological changes in marine fish at the species level [41]. Using the power of large numbers, community-wide advances in the phenology of fish larvae have been predicted based on quarterly surveys conducted over multiple decades [6], but no species-level predictions could be made. In this type of approach, quarterly sampling data have to be aggregated across many species and statistically interpolated to monthly averages before subjecting these interpolated values to a (linear) regression to arrive at an (extrapolated) prediction of phenological change of the community at a daily resolution. This complicated procedure highlights a mismatch between the level of observation and the desired level of inference. In species with highly variable abundances, such as herring, statistically interpolated data used to ‘fill’ missing temporal resolution will likely deviate considerably from catch values obtained with higher temporal sampling frequency. Our species-specific outcomes show how the interpretation of phenological shifts is complicated when moving from high-resolution daily sampling to weekly, biweekly, or monthly sampling frequencies. In our analysis, predicted catches declined a hundredfold with decreasing sampling frequency as the chance of sampling peak migration days was greatly reduced. Furthermore, visual inspection of smoother plots no longer revealed a clear phenological shift, or did not have ecologically relevant effect sizes with decreasing sampling frequency. Our results, therefore, stress the importance of long-term local monitoring surveys that can maintain high temporal resolution in improving our broader understanding of how different marine fish species respond to climate-driven changes by shifting their phenology of habitat use.

Changes in sea water temperature are known to be a major determinant of the distribution of small pelagic fish species such as clupeids [42, 43]. We found a clear signal of regional North Sea water temperatures on herring catches in the coastal Wadden Sea system. The North Sea is among the fastest warming large marine systems on the globe [44]. Furthermore, the Marsdiep tidal basin, in which the kom-fyke is situated, has experienced an approximate 1.5 *^◦^*C increase in temperature over the past 25 years [45]. The large positive effect of relatively cold North Sea water temperatures on weekly catches in herring aligns with our finding of a relative increase in herring catches earlier in the year with time. Warming and increased water temperature, therefore, provide the most credible explanation for the forward phenological shift observed in herring in the present study. From this perspective, changes in these local dynamics reflect larger-scale response patterns in fish populations observed in other regions of the world [5, 6]. Additionally, lunar illumination played a weak role in describing changes in the abundance of herring in our passive sampling gear. Lunar illumination can be important in the timing of migration movement as well as for prey availability and foraging success [46, 47].

Life-stage specific habitat utilization, such as juveniles using shallow coastal waters as nursery areas, is important for the life cycle closure and maintenance of large population sizes for many groups of marine species [48, 49, 50], including commercially important fish such as flatfish and clupeids [51, 52]. Here, we found life-stage specific changes in the coastal habitat use by North Sea herring. Changing suitability and use of such habitats due to climate-driven seawater warming can have potential broader, ecosystem-level consequences [53]. Warming sea waters greatly reduced the nursery function of the Wadden Sea for flatfish such as plaice (*Pleuronectes platessa*), dab (*Limanda limanda*), and flounder (*Platichthys flesus*), whereas conditions have improved for the more warm-tolerant sole (*Solea solea*) [20, 54, 55]. For pelagic species such as Atlantic herring, the long-term nursery function of the Wadden Sea was understudied. Our result that catches were dominated by small (5 to 10-cm) juvenile herring indicates that the Wadden Sea serves as a nursery for this species. This corroborates the recent results of Maathuis et al. [21] who reported large numbers of clupeids passing through the Marsdiep tidal inlet based on echosounder measurements. However, the long-term trend of decreasing herring abundance that we report here confirms a decline in this nursery function of this habitat [20]. Furthermore, in contrast to flatfish species [55], herring spawning stock biomass in the North Sea has also decreased in recent decades [56], suggesting that the decreasing nursery function of the Wadden Sea for herring is not directly compensated for by increased use of other nursery areas.

An important question that remains is how changes in phenology observed in the present study might impact the growth, development, and physiological status of sensitive life stages in marine fish. This is important because the rate of growth and development of juvenile fish is tightly linked to both ecological (competition, food availability), and physiological (thermal tolerance) processes that can have major impacts on population and community dynamics [57, 58]. For example, increasing temperatures in the Wadden Sea coincided with decreases in the in-situ abundance and, at the same temperatures, decreased individual growth in the laboratory in eelpout (*Zoarces viviparus*) [59]. Moreover, populations in temperate areas have been reported to be more sensitive to extirpation if juveniles are unable to build up enough reserves during times of abundant resources to get through resource-limited winters [60]. The timing of resource availability might shift under climate-driven warming and this has been suspected to lead to potential mismatches between the phenology of marine fish and the phenology of their prey resources [61] (but see [62, 63]). This widely studied but difficult to (dis)prove phenomenon is known as ‘phenological mismatch’ [64, 62], and the consequences of phenological mismatches have been studied most extensively in (migratory) birds [65]. The general consensus is that organisms at lower trophic levels are more likely to advance their phenology than organisms at higher trophic levels [66, 67], and that advancing phenology in resource availability negatively affects the growth and survival of individual offspring at higher trophic levels [68]. However, despite the negative impact on offspring at the individual level, no negative effect has been found on demographic measures at the population level [69]. The lack of a population level effect is thought to be due to density-dependent compensatory mechanisms, but this hypothesis remains untested. These studies from different systems show how understanding the general impacts of phenological shifts on fish populations requires wider research focus; where population level measures are linked to both fine-scale data on individual development as well as to resource availability. Making these types of links for marine fish will require improved integration of existing surveys and modelling methodologies in future studies. Combining local surveys with high temporal resolution of different trophic levels will play a key-role in this process by placing changes in marine fish phenology within a wider ecosystem context. This would enable studies to, for example, make fine-scale inferences on whether changing phenology of small juvenile fish follows changes in phytoplankton productivity; or whether changing phenology and body condition of small juvenile fish affect the prey selection and reproductive success of predators that are less likely to change their phenology, such as seabirds and marine mammals. In this way, local long-term high-resolution time series of fish catches can be valuable in defining general thresholds of climate-driven warming and aid in defining ecological indicators of system-level change [70].

## Acknowledgements

We would like to thank all the NIOZ staff, students, and volunteers involved in various ways in organizing and collecting the kom-fyke monitoring program over the past decades. We would like to thank Anita Koolhaas for her work on the fyke database and her help in the preparation of the dataset used in this study. Next, we would like to thank Luc de Monte, Dennis Mosk, Hidde Kresin, and Robert Twijnstra as the present fyke crew combining the role of boat drivers, managing the day-to-day operations, and emptying of the fyke. From the past fyke crew we would like to thank Sieme Gieles, Marco Kortenhoeven, Bas Wensveen, Suzanne Poiesz, and Henk van der Veer. We would also like to thank our dissection team, many of whom were part of the aforementioned fyke crews, for processing all the fish.

