## Supplementary Table 1, Supplementary Figure S1-13 for "Local reflects global: Life-stage dependent changes in the phenology of coastal habitat use by North Sea herring"

### Local reflects global: Life stage dependent changes in the phenology of herring moving between the North-and Wadden Sea.

**Table S1.** Six-point scale used to determine gonadal ripeness of individuals subsampled for dissection (N=480).

| Gonadal ripeness | Male description | Female description |
|---|---|---|
| 1 - Immature | Testes undeveloped or very thin line | Ovaries undeveloped or empty |
| 2 - Maturing / Recovering | Testes slightly thickened on one side | Eggs visible as small points in ovaries |
| 3 - Maturing / Recovering | Testes thickened and coloring white | Ovaries color orange, eggs well visible |
| 4 - Mature / prespawning | Testes thick and white, but no release of milt | Eggs swelling, but no release |
| 5 - Spawning | Milt secretion when pressure is applied | Eggs released when light pressure is applied |
| 6 - Spent | Testes flabby | Ovaries bloodshot and flabby. |

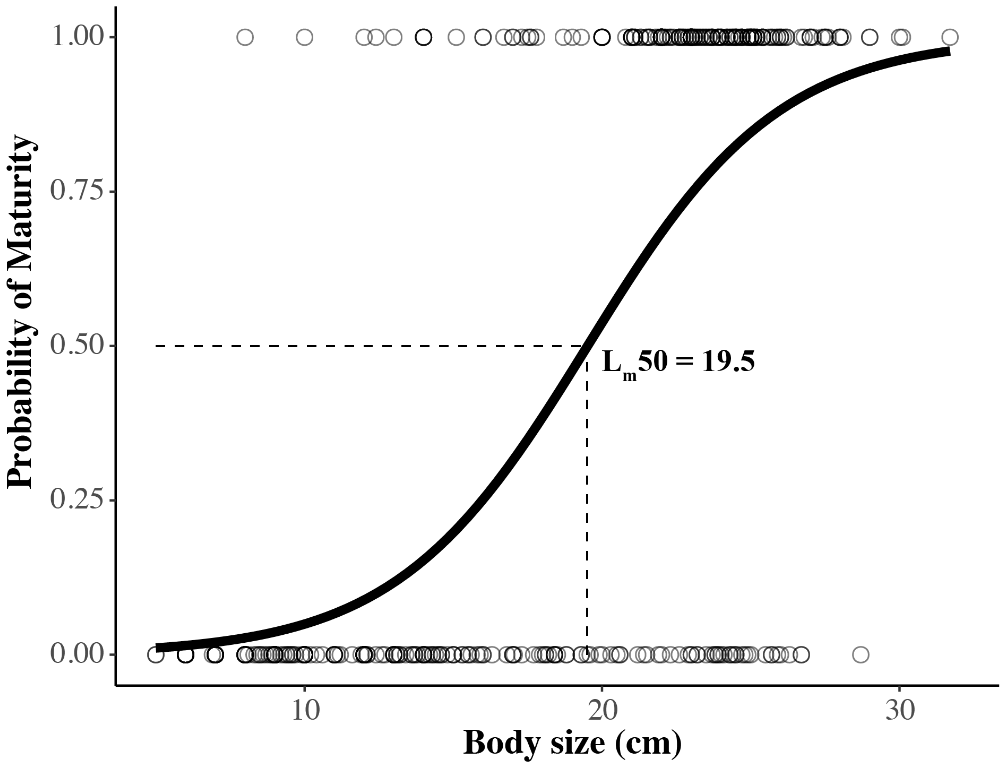


**Fig. S1.** Logistic regression to determine the size in cm at which 50% of all dissected individuals subsampled from the kom-fyke catches reach maturity ($L_m50$ = 19.5 cm, N=480). Individuals with Gonadal ripeness scores of 1 were classified as immature juveniles and individuals with gonadal ripeness scores > 1 as mature (see table S1).

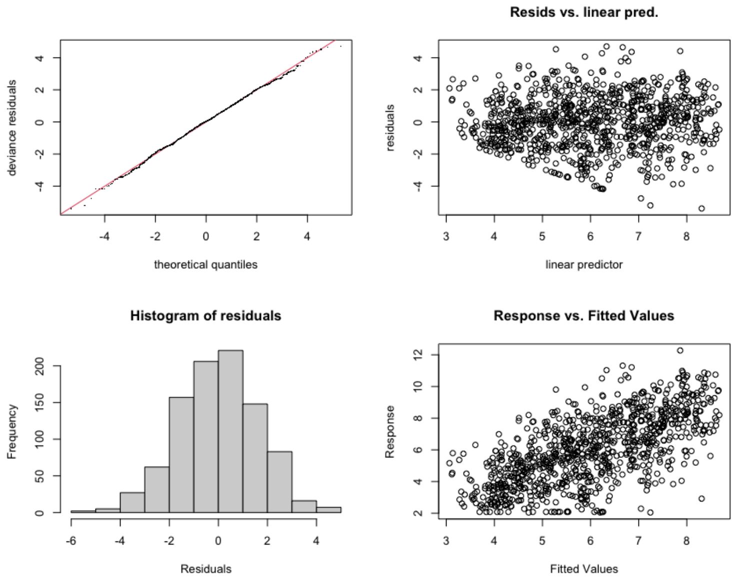


**Fig. S2.** Diagnostic plots for the additive model, clockwise: qqplot, residuals vs linear predictors, residuals histogram, and response vs fitted values.

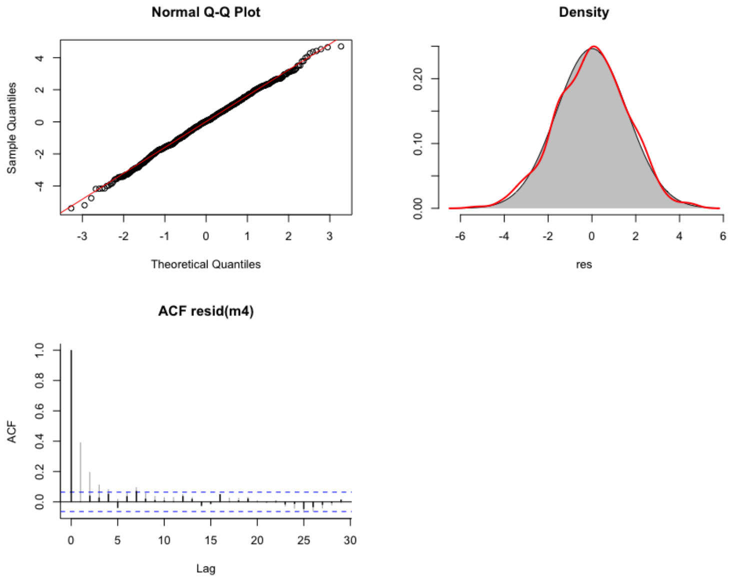


**Fig. S3.** Diagnositic plots for the additive model , clockwise: qqplot, residual density plot, and residual autocorrelation.

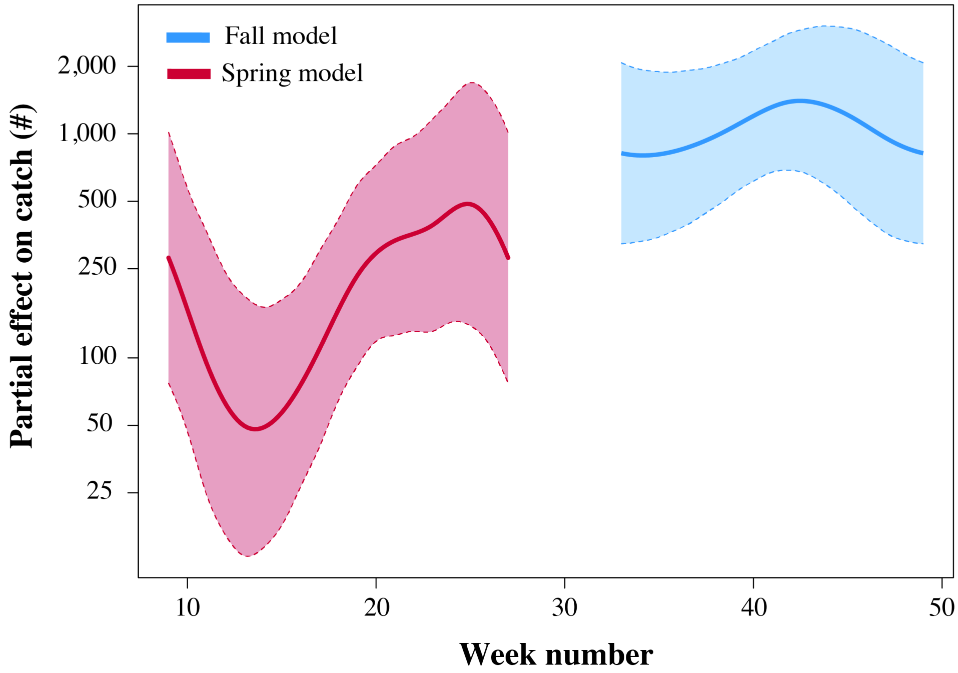


**Fig. S4.** Partial interaction effect of week number on weekly standardized herring catches in the spring data (red line) and fall data (blue line) models used to compare against full data model in Fig.3a of the manuscript. Shaded area represents the pointwise 95% CI.

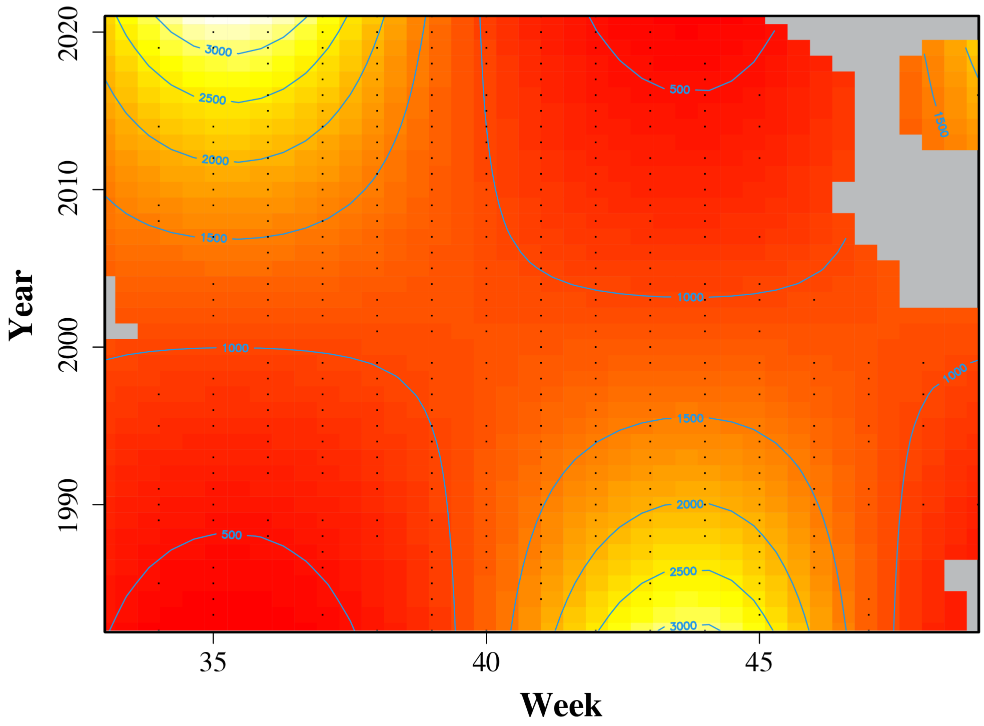


**Fig. S5.** Partial interaction effect of the week and year on weekly standardized herring catches in the fall data model used to compare against the full data model in Fig. 3b of the manuscript.

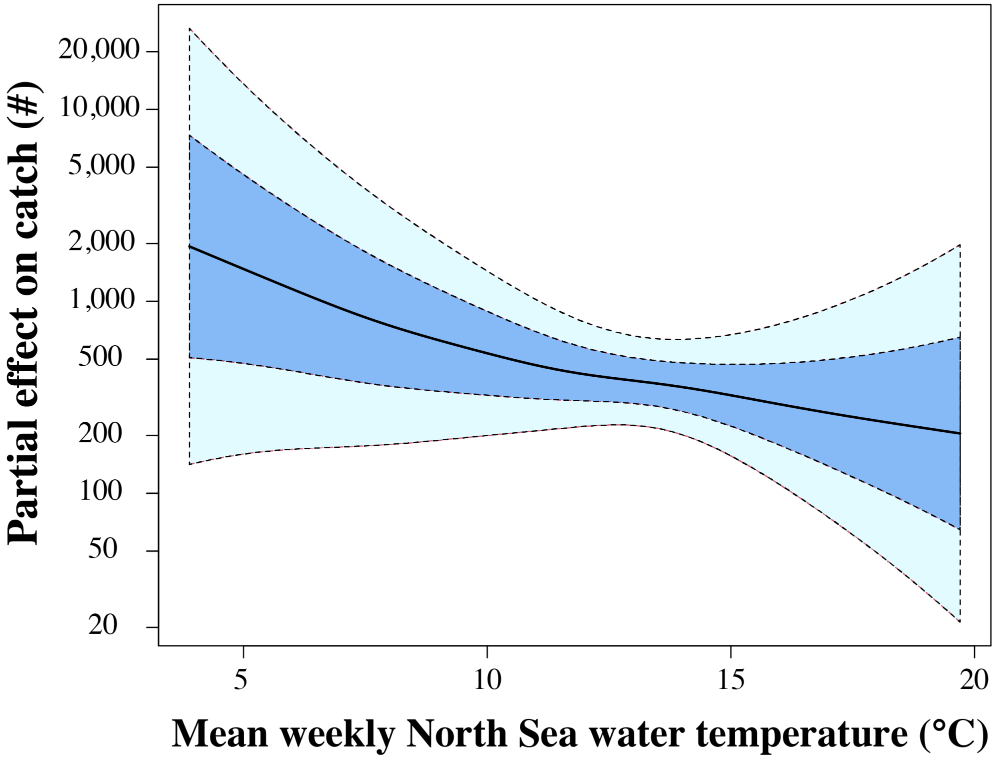


**Fig. S6.** Partial effect of North Sea water temperatures on weekly standardized herring catches in the full data model. Shaded dark blue area represents the point-wise standard errors, and the shaded light blue area the pointwise 95% CI.

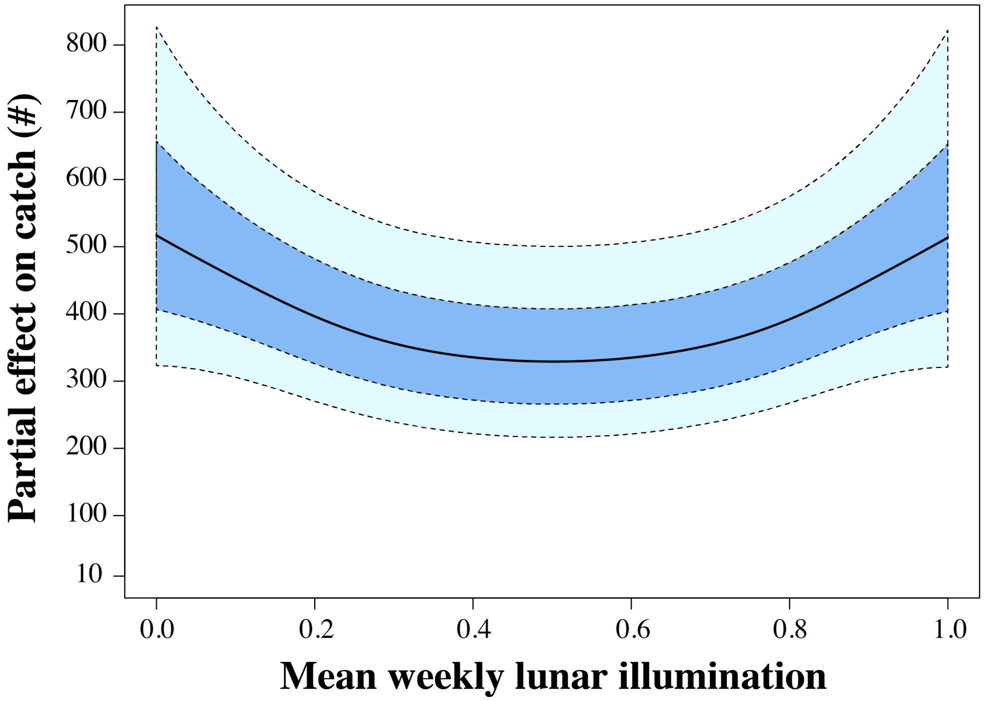


**Fig. S7.** Partial effect of lunar illumination on weekly standardized herring catches in the full data model.Shaded dark blue area represents the point-wise standard errors, and the shaded light blue area the pointwise 95% CI.

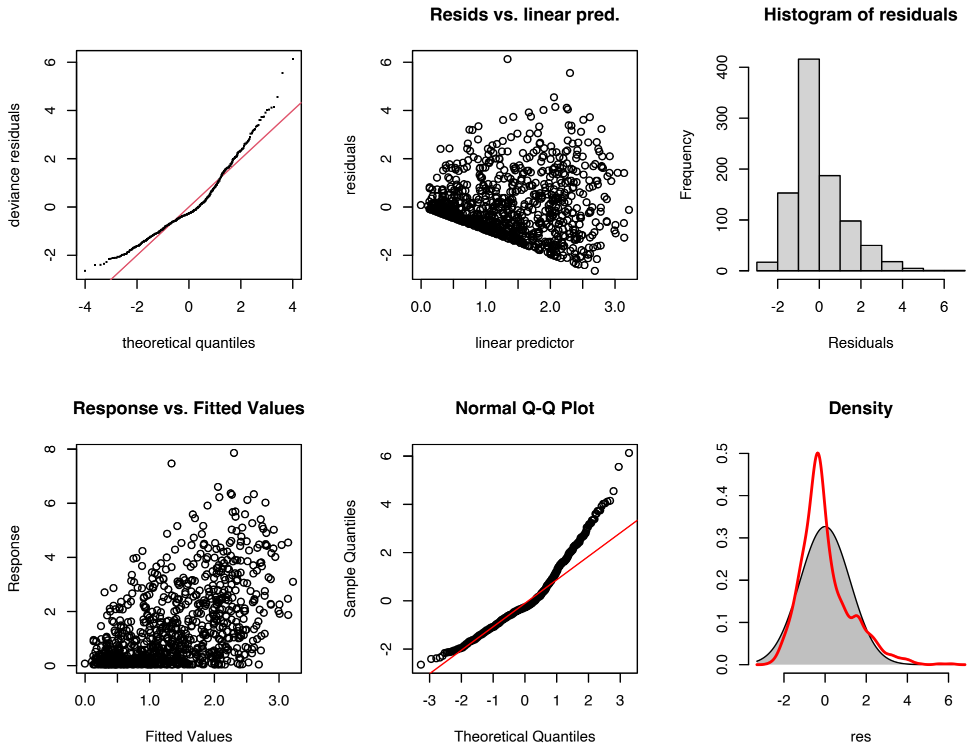


**Fig. S8.** Diagnostic plots for the weekly sampled additive model, clockwise: qqplot, residuals vs linear predictors, residuals histogram, and response vs fitted values.

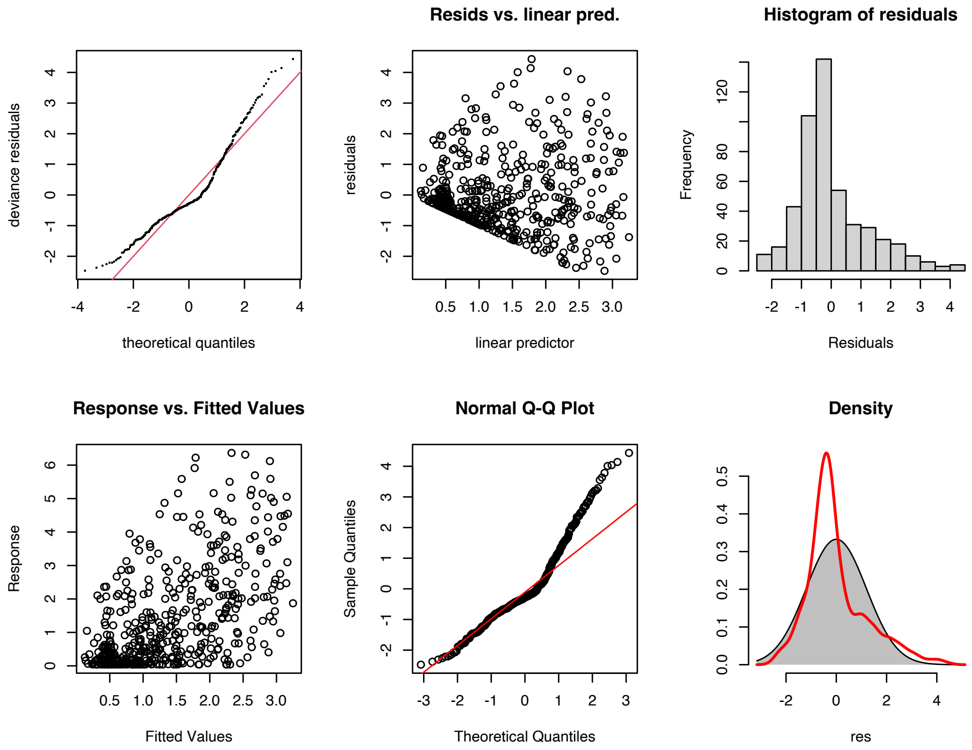


**Fig. S9.** Diagnostic plots for the biweekly sampled additive model, clockwise: qqplot, residuals vs linear predictors, residuals histogram, and response vs fitted values.

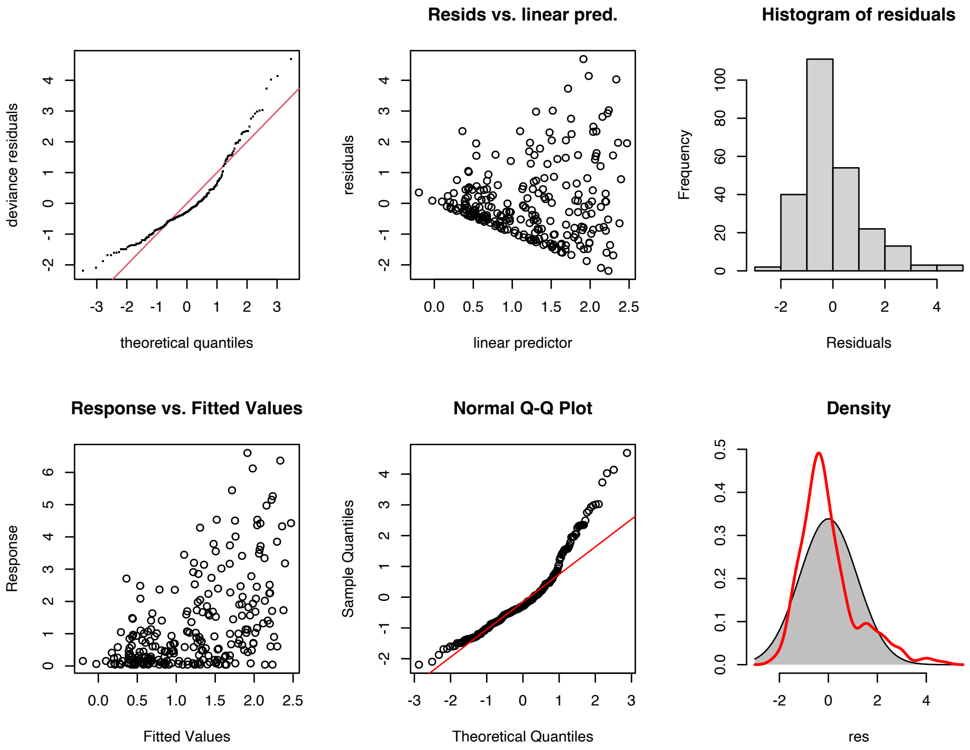


**Fig. S10.** Diagnostic plots for the monthly sampled additive model, clockwise: qqplot, residuals vs linear predictors, residuals histogram, and response vs fitted values.

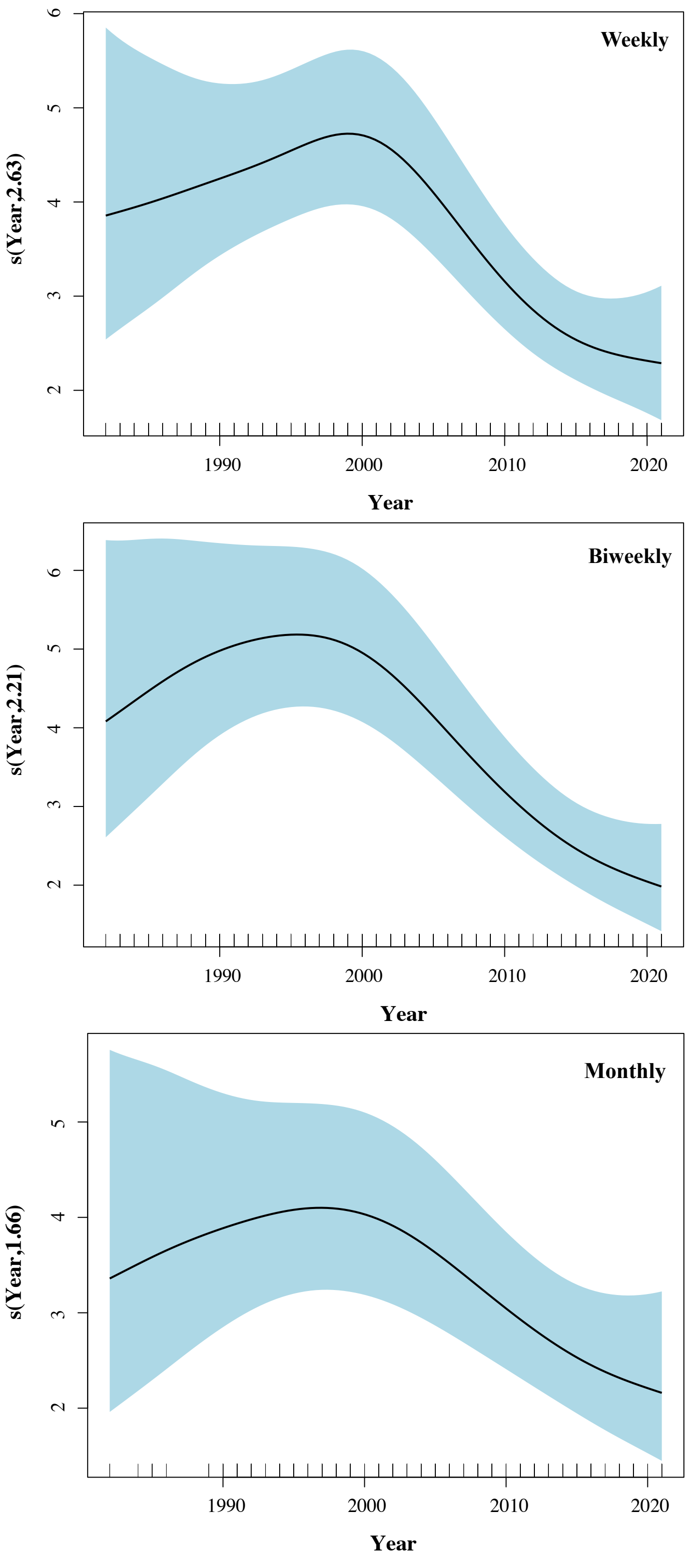


**Fig. S11.** Partial effect of year on standardized herring catches when sampling once per week, biweekly, or month, respectively.

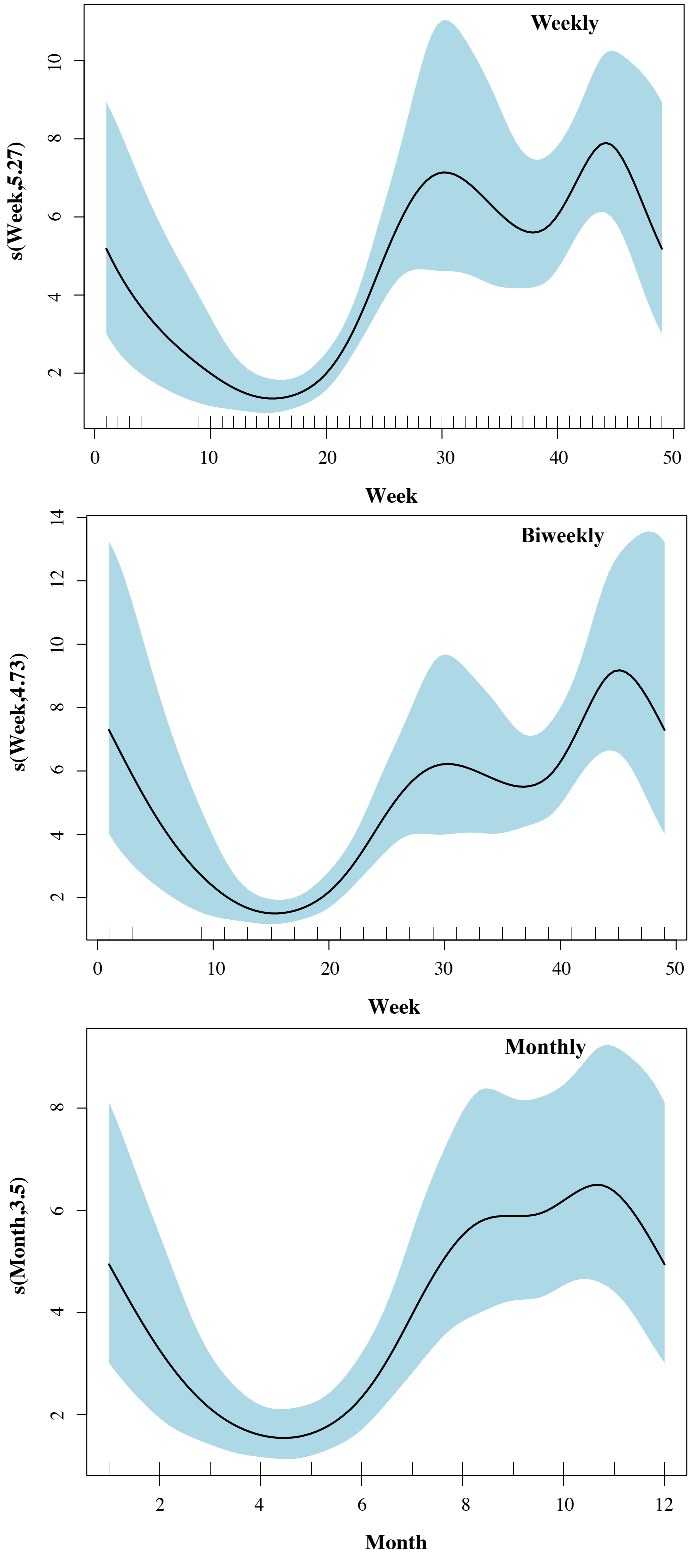


**Fig. S12.** Partial effect of season on standardized herring catches when sampling once per week, biweekly, or month, respectively.

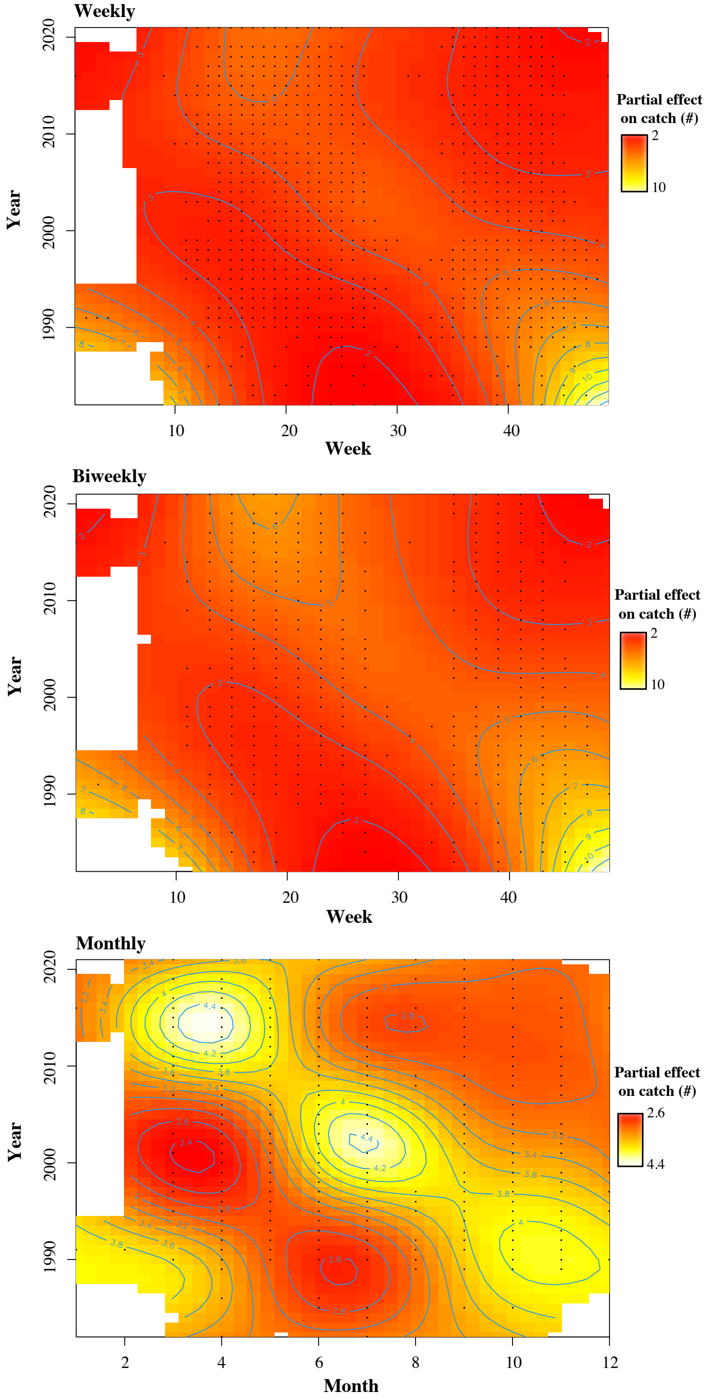


**Fig. S13.** Partial interaction effect of season and year on standardized herring catches when sampling once per week, biweekly, or month, respectively.
